# The neoantigen landscape of mycosis fungoides

**DOI:** 10.1101/2020.01.22.915280

**Authors:** A Sivanand, D Hennessey, A Iyer, S O’Keefe, P Surmanowicz, G Vaid, R Gniadecki

## Abstract

**Background:** Mycosis fungoides (MF), the most common type of cutaneous T-cell lymphoma, has a dismal prognosis in advanced stages. Treatments for advanced disease are mostly palliative and MF remains incurable. Although MF is a known immunogenic neoplasm, immunotherapies such as interferons and the immune checkpoint inhibitors yield inconsistent results. Since the number, HLA-binding strength and subclonality of neoantigens are correlated with the therapeutic responses, we aimed here to characterize the landscape of neoantigens in MF.

**Methods:** We conducted whole exome and whole transcriptome sequencing of 24 MF samples (16 plaque, 8 tumour) from 13 patients. Bioinformatic pipelines (Mutect2, OptiType, MuPeXi) were used for *in silico* mutation calling, HLA typing, and neoantigen prediction. Phylogenetic analysis was used to subdivide the malignant cell population into stem and clades (subclones). Clonality of neaontigens was determined by matching neoantigens to the stem and clades of the phylogenetic tree of each MF sample.

**Results:** MF has a high mutational load (median 3217 non synonymous mutations), resulting in a significant number of total neoantigens (median 1309 per sample) and high-affinity neoantigens (median 328). In stage I disease most neoantigens were clonal but with progression to stage II, subclonal neoantigens comprised >50% of the total. There was very little overlap in neoantigens across patients or between different lesions on the same patient, indicating a high degree of genetic heterogeneity.

**Conclusions:** Analysis of the neoantigen landscape of MF revealed a very high neoantigen load and thus a significant immunogenic potential of this lymphoma. However, neoantigenic heterogeneity and significant subclonality might limit the efficacy of immunotherapy. We hypothesize that neoantigen number and subclonality might be useful biomarkers determining sensitivity to immunotherapeutic strategies.

## Introduction

Mycosis fungoides (MF) is the most common type of cutaneous T-cell lymphoma (CTCL) (1) that develops from clonotypically diverse malignant T-cell precursors seeding the skin (2,3). Prognosis in the early stages (T1-T2, patches and plaques) is excellent, however the development of tumours (T3) or erythroderma (T4) is associated with a significant decrease in survival (4,5). Despite intensive research, MF remains incurable and treatments for advanced disease are mostly palliative (5).

There is robust evidence that MF is an immunogenic tumour and that the immune system is an essential factor limiting its progression (reviewed in ref. (6)). It has been well documented that iatrogenic immunosuppression causes a catastrophic dissemination of MF (7,8). Many current therapies (interferons, imiquimod, extracorporeal photopheresis and allogeneic stem cell transplant) are considered to act primarily via stimulation of the antitumour immunity (9–12). However, the experience with immune checkpoint inhibitors has been disappointing in MF (6). The literature comprising approximately 50 cases of MF treated with various immune checkpoint inhibitors reports response rates ranging from 9% to 56% with only a few documented complete remissions (13–17). Of the few anticancer vaccine studies in CTCL, response rates have ranged from 33% to 50% (18–20). Those rather discouraging results are surprising in view of the fact that MF is a mutationally rich tumour with a mutation load in the range of 500-4,500 somatic mutations/genome (21). The number of mutations is usually correlated with the number of neoantigens and consequently the immunogenicity of the cancer, which is predictive for immune checkpoint inhibitor efficacy (22–24).

It has recently been suggested that in addition to mutational load and the number of neoantigens, tumour heterogeneity has a major impact on the ability of the host immune system to mount an effective antitumour defense. Neoantigens can be classified as clonal (present on all cancer cells) or subclonal (present only on a subset (subclones) of cancer cells) (22). A high clonal neoantigen burden, for instance in malignant melanoma, favours effective immune surveillance, response to immunotherapy and significantly prolonged survival (22). In contrast, a tumour with a branched subclonal structure will be poorly recognized by the immune system, even if the mutation load is high, as documented for some immunotherapy-resistant tumours such as glioblastomas (25).

To better understand the potential for immunotherapeutic approaches in MF we studied the landscape of neoantigen expression in this malignancy. Using whole transcriptome and whole exome sequencing, we determined the pattern of neoantigens in early lesions of patches and plaques and compared them to those of clinically advanced disease. We show that disease progression is correlated with an increase in mutational load and the number of neoantigens. However, advanced lesions of MF exhibit a high proportion of subclonal neoantigens which may limit the efficacy of immunotherapies.

## Methods

### Materials, sequencing, datasets

Institutional ethics approval was obtained under the application HREBA.CC-16-0820-REN1. We performed whole exome sequencing (WES) and whole transcriptome sequencing (WTS) of 24 MF samples (16 plaque, 8 tumour) and matched peripheral blood mononuclear cell (PBMC) in 13 patients (patient characteristics in **Supplementary Table S1**). DNA and RNA sequencing libraries were prepared from tumour cell clusters microdissected from skin biopsies using laser capture microdissection and sequenced as described previously (21,26) (Figure 1). Additional datasets comprised sequencing data published by McGirt et al. (5 whole genome sequences (WGS) from 5 patients with MF) (27) and by Choi et al. (31 WES from 31 patients with Sézary syndrome) (28).

**Figure 1:**
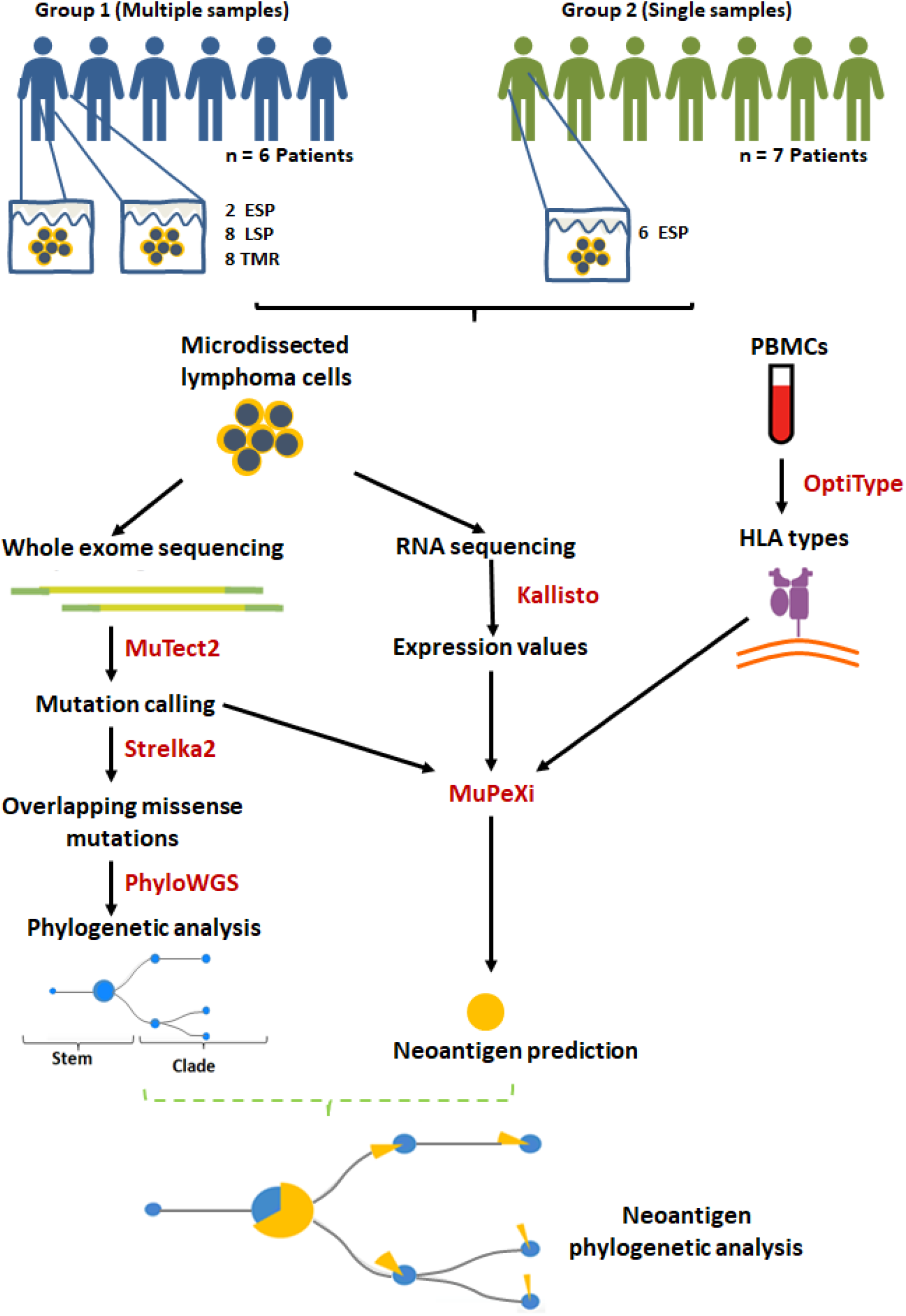
Summary of methods and study design. Biopsies of lesional skin and blood were obtained. 13 MF patients were divided into group 1 (multiple samples) and 2 (single samples) according to the number of biopsies contributed. The lesions were categorized according to the clinical stage and the morphology of the lesion: ESP (early stage plaques, i.e. MF plaques in stage I), LSP and TMR (respectively, late stage plaques and tumours biopsied from patients in stage ≥IIB). MuPeXi was used to predict neoantigens. For clonality analysis we used mutation data obtained from MuTect2 and Strelka2, as described previously (21). Predicted neoantigens were mapped to the clades and stems of the phylogenetic trees constructed using PhyloWGS (21).

### Identification of neoantigens

Bioinformatics analysis involved a series of pipelines shown in Figure 1. GATK (v4.0.10) best practices guidelines (29) were used to process the initial WES fastq files. Reads were aligned to the hg38 reference genome. MuTect2 (v2.1) was used for variant calling to identify missense and indel mutations. OptiType (v1.3.1) (30) was used with default settings to predict class I human leukocyte antigen (HLA) types from WES of PBMC for the corresponding samples. Kallisto (v0.45.0) (31) was used to process the raw RNA fastq files with the bootstrapping function set to 500 to obtain the variance and expression level. The outputs of these pipelines (.vcf files from MuTect2, HLA types from Optitype and .tsv files from Kallisto) were imported into MuPeXi (v1.2) (32) to predict neoantigenic peptides (8-11 amino acids long). NetMHCpan 4.0 (33) (incorporated in MuPeXi pipeline) was used to predict peptide binding affinities to up to 6 patient-specific HLA types.

### Neoantigen filtering

We will refer to the raw output of prediction software as ‘putative neoantigens’ and the result once filtering criteria is applied as ‘filtered neoantigens’. Our filtering criteria included: (1) Mutant peptide binding strength, defined as eluted ligand (EL) likelihood percentile rank ≦0.5%, (2) RNA expression level >0.1 transcripts per million (TPM) (34), (3) Top peptide, applied last to group all predictions arising from the same mutation (chromosome and genomic position) and select the peptide with the lowest binding strength. While all peptides <0.5% rank are generally considered to be strong binders (35), we further divided these into high strength binders (<0.05%rank), intermediate strength binders (0.05≧%rank<0.15) and low strength binders (0.15≧%rank≦0.5).

### Mutant peptide characterization

To further characterize mutant peptides, we identified the most frequently overlapping peptides between samples. We then used the mutant peptide sequence to search the IEDB database (36) for homologous peptides that were known immune epitopes. We searched for exact matches and if none were found, we reduced the threshold to blast >90%. If a known epitope was found, we further searched the Uniprot database (37) for details of the gene encoding the protein, and the protein function.

### Neoantigen clonality analysis

For phylogenetic analysis, Strelka2 (v2.9.10) (38) was used for mutation calling to identify missense mutations that overlapped with those called by MuTect2. TitanCNA (39) was used to predict copy number aberrations (CNA). Default parameters were used except for alphaK which was changed to 2,500 as recommended for WES data. PhyloWGS (v1.0-rc2) (40) was used to build phylogenetic trees by clustering missense mutations using CNA. The stem and clade mutations producing neoantigens were then highlighted on the phylogenetic trees to determine the clonality of the neoantigens.

### Data visualization

Visual data representations were created using the R package beeswarm (41), GraphPadPrism (v8.3.0) (42), jvenn (43), Venn Diagram Tool (44) PhyloWGS (40) and Microsoft Excel.

### Data availability

The de-identified whole exome sequencing data have been deposited to the database of Genotypes and Phenotypes (dbGAP). The accession code to this data is phs001877.v1.p1.

## Results

### Tumour mutation burden in MF is dominated by frameshift mutations

Early-stage MF (IA-IIA) is characterized by thin cutaneous lesions of patches and plaques (T1-T2). The emergence of tumours (T3) heralds progression to the advanced stage IIB. It is important to note that most advanced-stage patients may exhibit plaques persisting from the early stages in addition to the stage-defining tumours. To capture the impact of disease stage on mutation burden and neoantigen expression we classified biopsies into the following categories: early stage plaques (ESP), i.e. the lesions T1 and T2 (patches or plaques) obtained from patients in stage IA-IB, and late stage plaques (LSP) and matched tumours (TMR) from patients in a clinical stage ≥ IIB. In those lesions, we determined tumour mutation burden (TMB) defined as the number of non-synonymous mutations producing neoantigens. The median TMB was 3,217 mutations per sample, or 35 mutations/kB comprising primarily of frameshift mutations (70.3%), in-frame missense mutations (28.4%), insertions (1.1%) and deletions (0.2%) (Figure 2). The median TMB in ESP was 2,455 (range 1,440-7,198), and its upper range increased in LSP (median 5014, range 890-8,697) and in TMR (median 2,697; range 1306-8,722).

**Figure 2:**
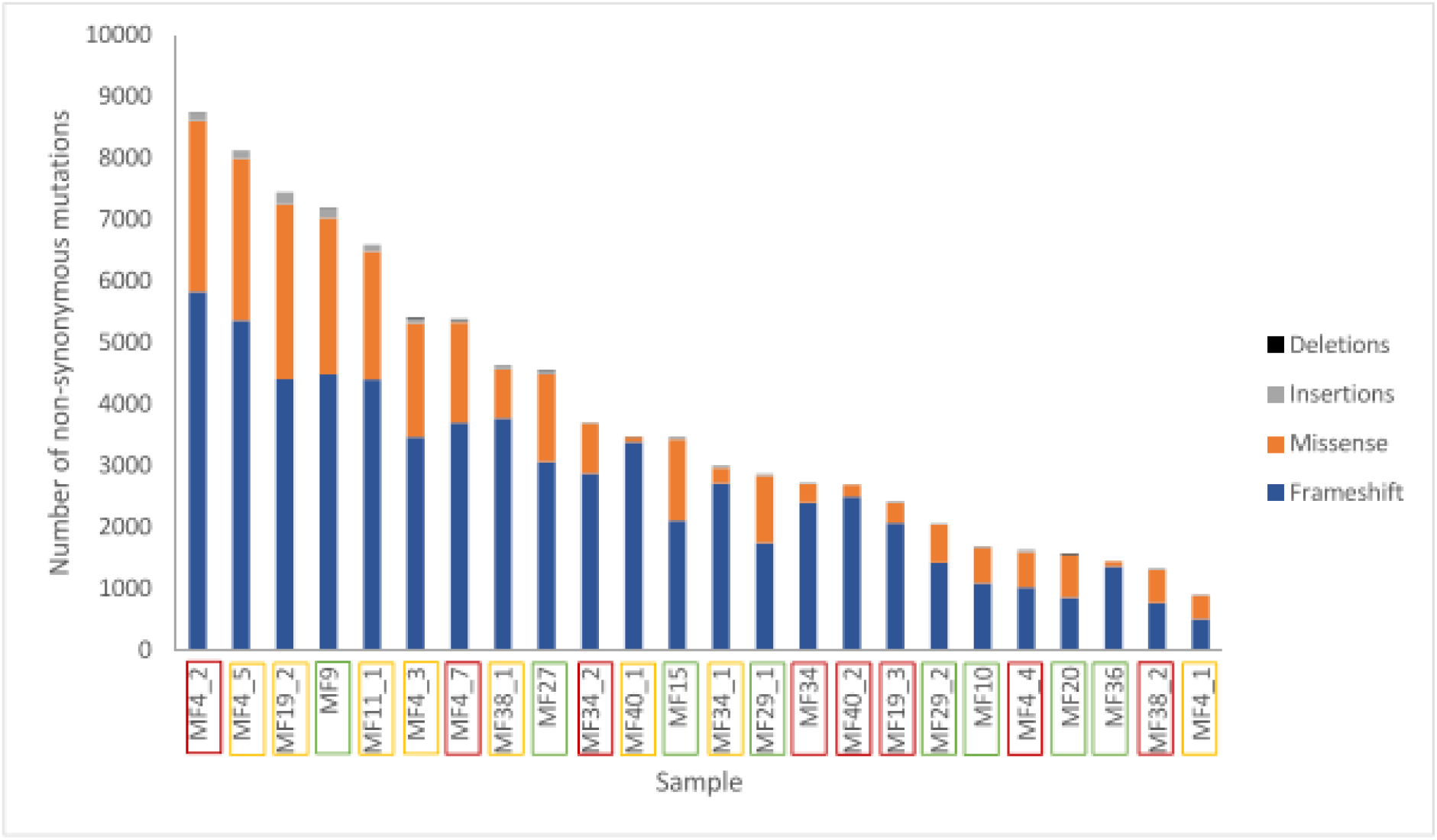
Tumour mutation burden. Samples are arranged in descending order of TMB. Frameshift mutations comprise the majority of non-synonymous mutations. Sample names are enclosed in boxes with colours corresponding to the lesion type - early stage plaque (green), late stage plaque (yellow) and tumour (red).

### Increase in neoantigen load during disease progression

When examined by lesion type, lesions in patients with advanced disease had a greater number of putative neoantigens compared to early stage plaques (LSP - 27,179,348, TMR - 19,647,017 vs ESP - 15,645,072) (Figure 3A). There was no difference in median binding strength between ESP (median 56%), LSP (median 57%) and TMR (median 58%).

**Figure 3.**
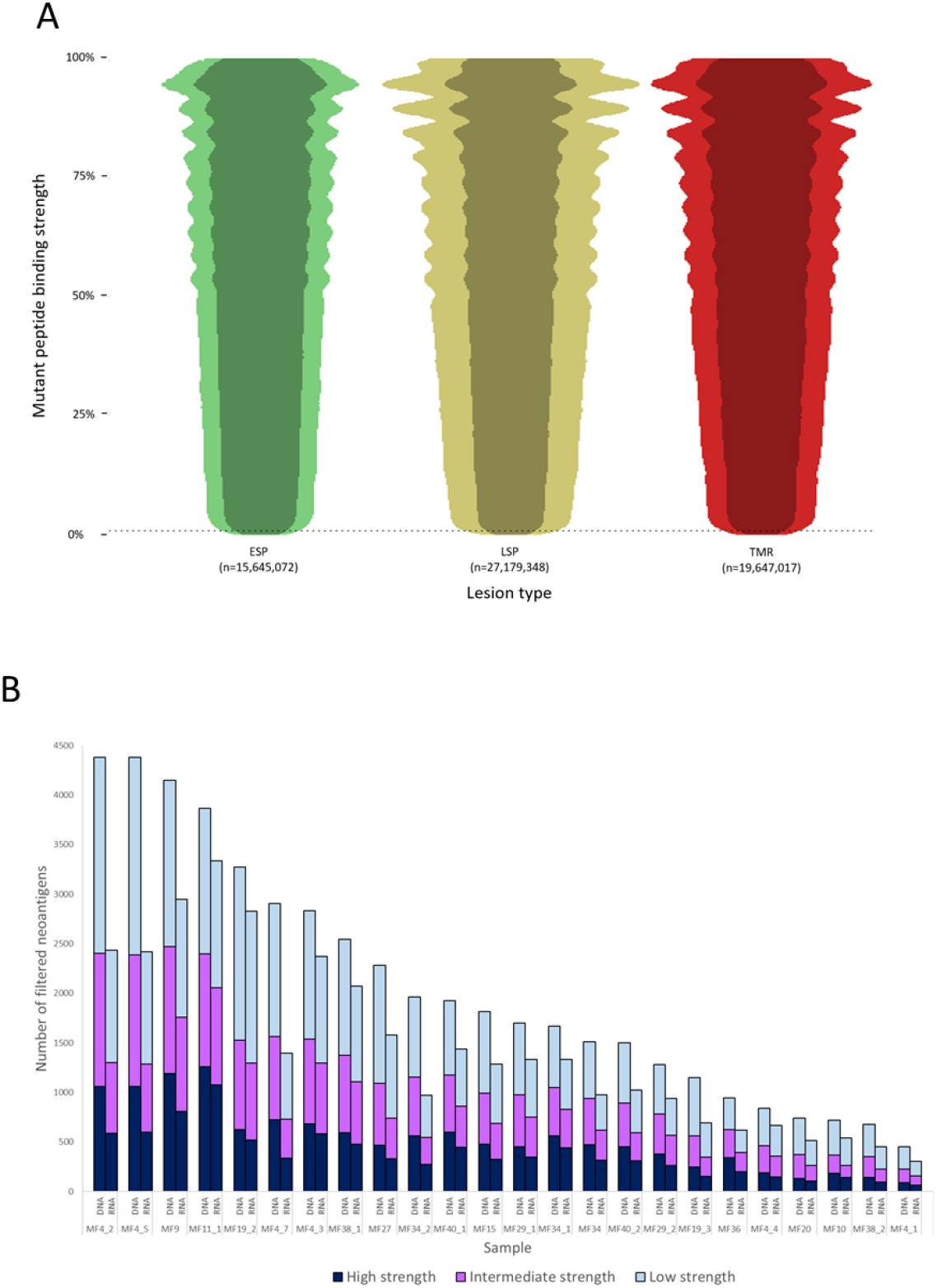
A: A beeswarm plot representation of putative neoantigens prior to filtering. Due to the extensive size of the dataset, a random 1% of all data points were plotted to demonstrate the overall distribution and density of the data. The vertical axis shows mutant peptide binding strength as a percentile rank, with lower values representing increasingly strong binding peptides to HLA types. 0.5% rank (dashed line) represents the commonly used cutoff below which peptides are considered strong enough binders to be neoantigens. The width of each plot is proportional to the number of neoantigens at each binding strength. Overall, ESP lesions had fewer neoantigens compared to LSP and TMR. The darker shade within each plot represents the neoantigens expressed in RNA (TPM>0.1). **B: Neoantigen load before and after applying the RNA filter.** For each sample, the “DNA” column has all filters applied with the exception of the RNA filter. The median number of filtered neoantigens per sample was 1,309. The “RNA” column has all filters including the RNA filter (expression >0.1 TPM) applied. On average 70% of predictions were expressed in RNA.

Filtering putative neoantigens is necessary to narrow down epitopes that are most likely expressed in patients. When we applied all filters (“RNA” column in Figure 3B), an average of 70% of predicted neoantigens were expressed at the RNA level (a median of 1,309 neoantigens per sample). A median of 328 were high strength binders (<0.05%rank), 376 were intermediate strength binders (0.05≧%rank<0.15) and 540 were low strength binders (0.15≧%rank≦0.5).

We further compared the association between tumour mutation burden and the filtered neoantigen load (Figure 4A), which showed a strong positive linear relationship (r=0.92). The tumour mutation burden also demonstrated a positive linear relationship with the number of high strength neoantigens (r=0.81). The tumour mutation burden, filtered neoantigen load and number of high strength neoantigens are summarized in Figure 4B.

**Figure 4.**
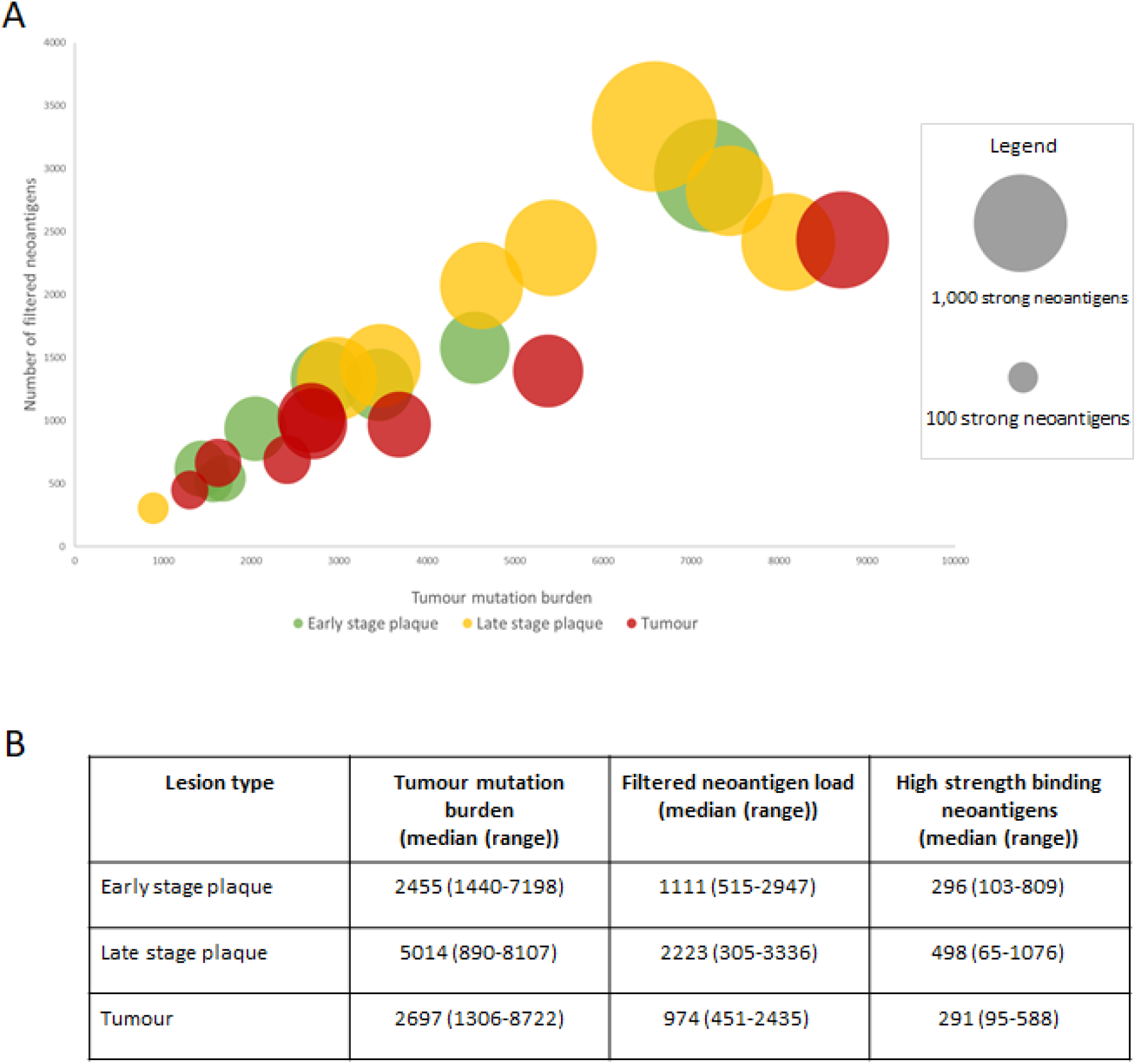
A: Tumour mutation burden and filtered neoantigen load. **A:** A strong positive linear association (r=0.92) was observed between tumour mutation burden and filtered neoantigen load. Each bubble represents a single sample, with its size proportional to the number of high strength neoantigens (<0.05%rank). A positive linear association was also observed between tumour mutation burden and the high strength neoantigen load (r=0.81). **B:** Mutations and neoantigen numbers by lesion type.

Comparing our data to the two previous CTCL studies of McGirt et al (27) and Choi et al (28), we found that our dataset had a much higher neoantigen count (total 54,073,746 vs 615,761 in Choi et al (28) or 135,042 in McGirt et al (27), **Figure S1)**.

### Increase in proportion of subclonal neoantigens in advanced MF

To determine the subclonality of the neoantigens we first constructed phylogenetic trees showing the subclonal architecture of MF, as described previously (21). This allowed us to map neoantigens to the stem and clades, the latter representing the subclonal neoantigens (Figure 5A). This analysis demonstrated an increasing branching with a higher proportion of clade neoantigens in advanced lesions, as demonstrated in LSP (median 62% clade neoantigens) and TMR (median 70%), compared to ESP (median 39%) (Figures 5A & 5B).

**Figure 5.**
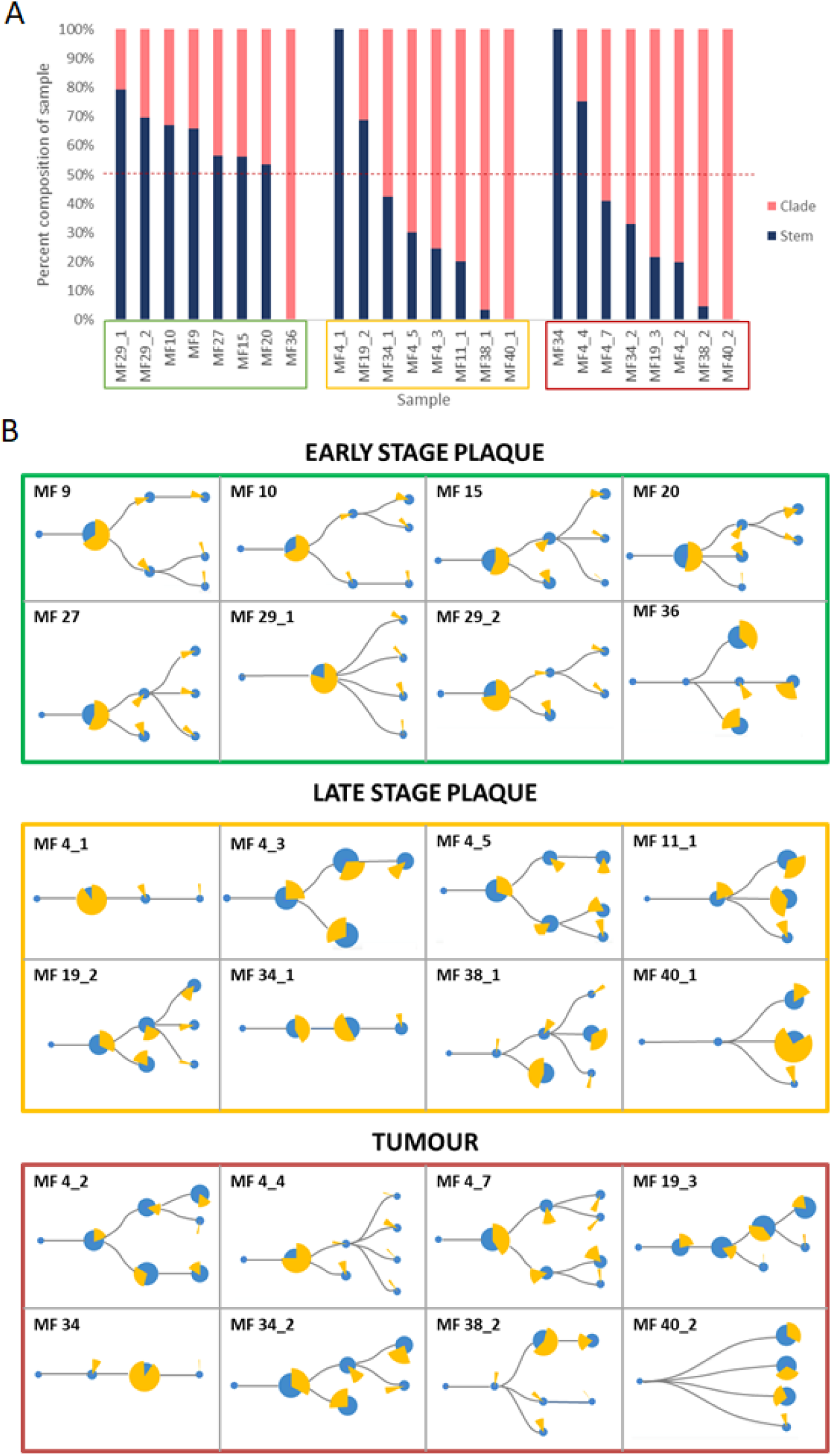
A: Proportion of stem and clade missense mutations producing putative neoantigens. **A:** Sample codes are enclosed in boxes with colours corresponding to the lesion type - early stage plaque (green), late stage plaque (yellow) and tumour (red). The dashed red line represents the 50% mark that distinguishes whether the majority of the sample is composed of stem or clade neoantigens. Early stage plaques have a greater proportion of stem mutations producing neoantigens compared to late stage plaques and tumours where more clade mutations produce neoantigens. **B:** Phylogenetic trees with putative neoantigen analysis. The size of the blue circles represents the proportion of missense mutations that comprise each node. ‘Stem’ nodes are those present prior to branching which then produces ‘clade’ nodes. The yellow pie chart in whole represents all neoantigens from the sample. Each slice of the pie chart represents the proportion of neoantigens originating from a node. With advancing disease stage, a greater proportion of neoantigens originate from clade mutations.

### Neoantigen overlap and peptide identity

We examined the overlap in filtered neoantigens by lesion type (Figure 6A) and within the same patient sampled longitudinally (Figure 6B). The overlap between late stage plaques (LSP) and tumours (TMR) was greater than between early stage plaques (ESP) and either lesion type. This was expected, as we separated advanced disease into LSP and TMR for analysis in our study. We also examined neoantigens from one patient from whom multiple, longitudinal samples were obtained. Among 6 samples (3 TMR, 3 LSP) obtained at 3 timepoints (0, 9 and 10 months respectively), we found no overlap in filtered neoantigens (Figure 6B). Most peptides were unique to each plaque or tumour site, further underscoring the predominantly subclonal structure of neoantigens in advanced disease. Finally, we examined the overlap of filtered neoantigens across samples (Figure 6C). No neoantigens were common to all samples, and the most common neoantigen was present in half of the 24 samples. Overlapping neoantigens were mostly present in late stage plaques and tumours. This is likely because advanced disease samples produced more neoantigens overall, increasing the likelihood of overlap.

**Figure 6.**
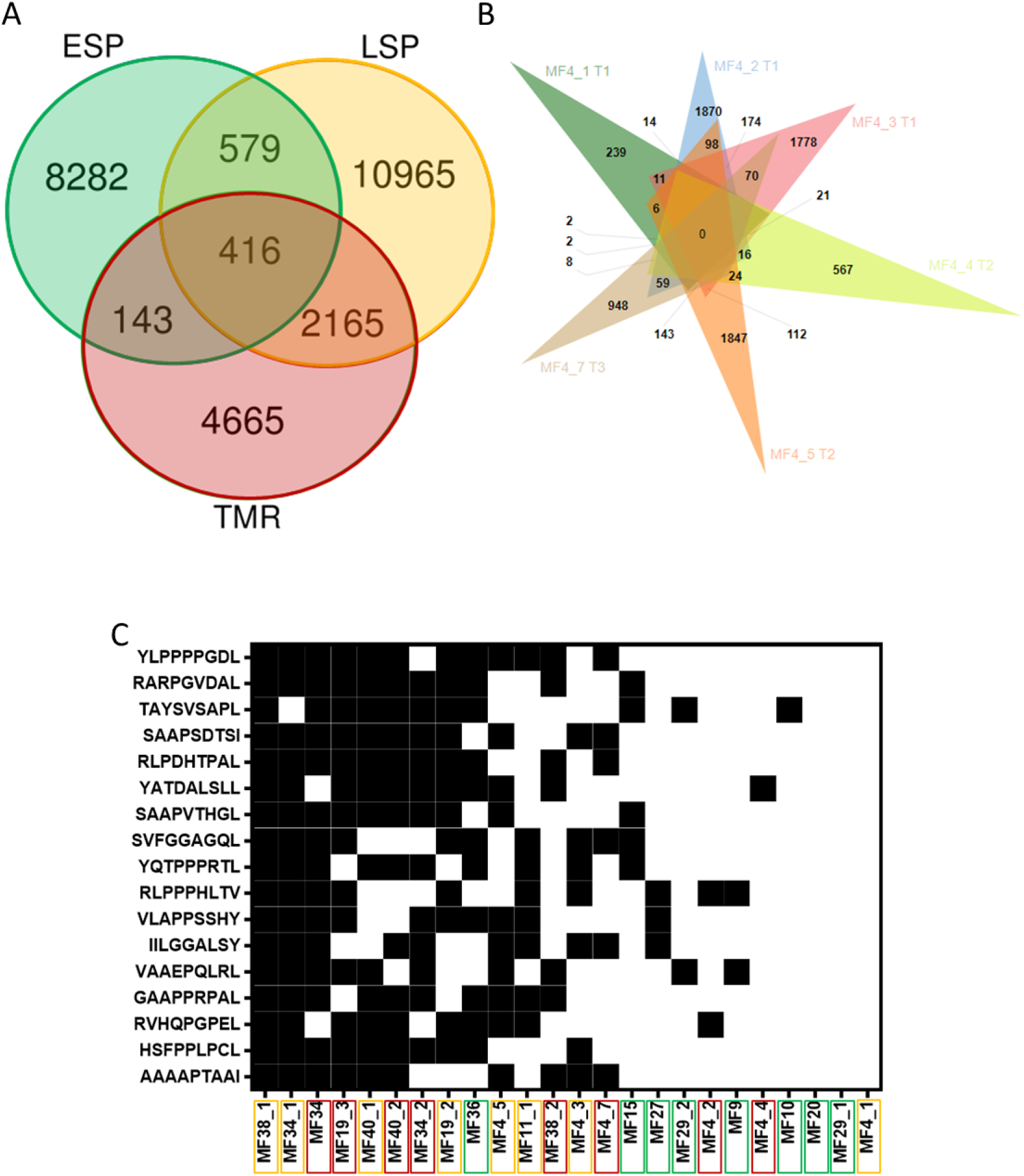
A: Intraindividual and interindividual overlap of neoantigens. **A:** Each lesion type comprises 8 samples, of which only unique peptides are included. The greatest overlap in filtered neoantigens is between plaques (LSP) and tumours (TMR). Early stage plaques (ESP) are also shown. **B:** Venn diagram of filtered neoantigens from 6 samples obtained from one patient. Each sample name is accompanied by the time point the biopsy was obtained (initial biopsy at T1, T2 at 9 months after T1 and T3 10 months after T1). There is no overlap in peptides between all lesions, and the predominant exclusivity of peptides to their individual sites indicates the highly branched nature of the tumour. **C:** Filtered neoantigens predicted in 10 or more samples out of the total 24 samples. Black indicates the presence of the peptide in the sample and white indicates the absence. Peptides are arranged from highest frequency (top) to lowest frequency (bottom). Sample names are arranged in order of those with the most overlapping neoantigens (left) to the least overlapping neoantigens (right). Sample names are enclosed in boxes with colours corresponding to the lesion type - early stage plaque (green), late stage plaque (yellow) and tumour (red). Overlapping neoantigens are mostly in the advanced stage disease samples (late stage plaque and tumour) clustered on the left.

Using the neoantigens we found, we searched IEDB for closely related peptides from humans or human pathogens (Table 1). These known immune epitopes have been tested in experimental assays and are likely to elicit immunogenic responses in humans. We included epitopes tested in T-cell, B-cell and MHC ligand assays and did not require assays to be positive. Only 2 neoantigens were positive in T-cell assays.

**Table 1:**
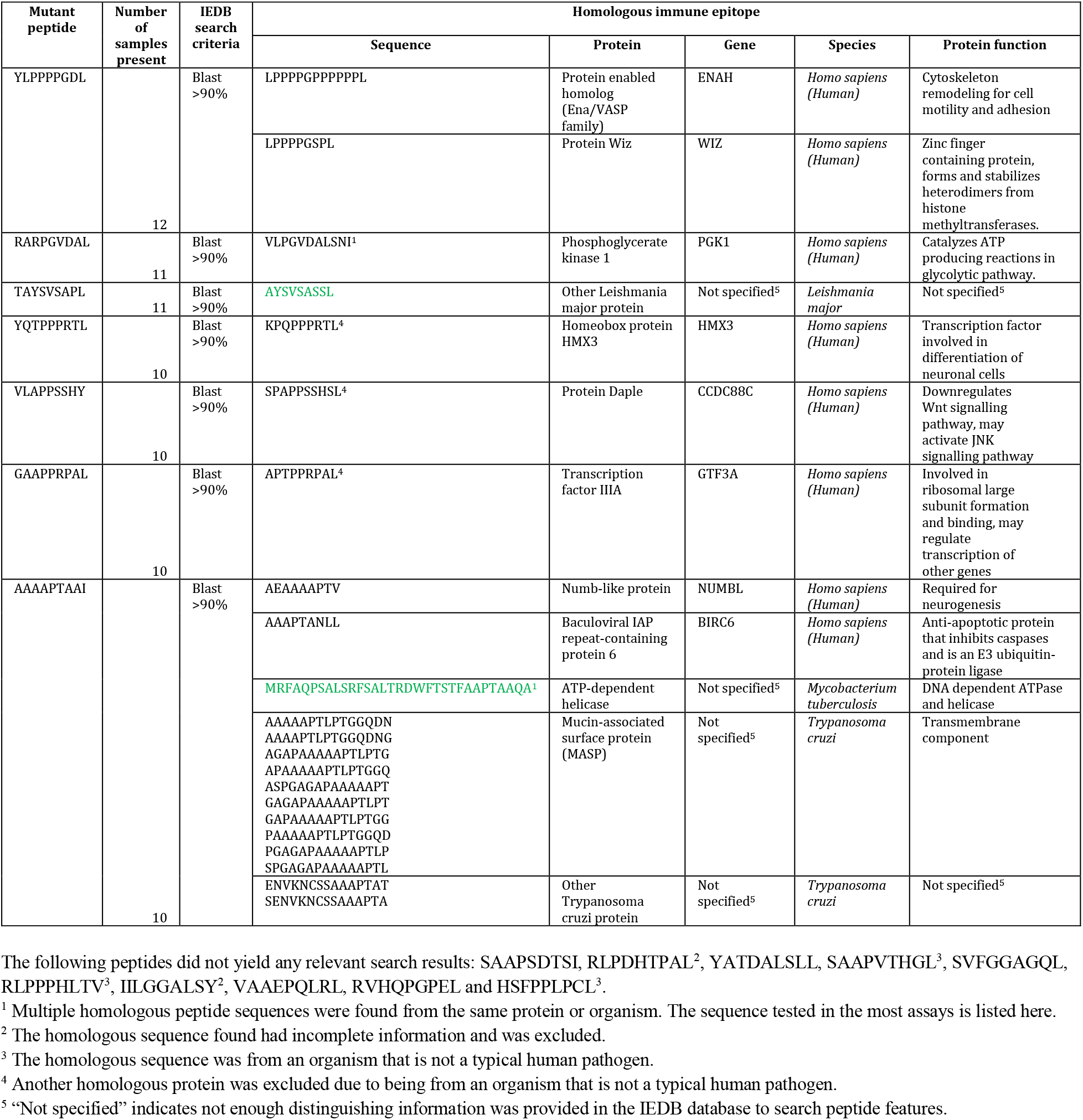
Homologous immune epitopes of filtered neoantigens. Only epitopes from humans or human pathogens were included. Included are epitopes tested in T-cell, B-cell and MHC ligand assays. There was no requirement that assays be positive. Peptides highlighted in green were positive in T-cell assays.

## Discussion

We have previously demonstrated that as MF progresses from early to advanced stages, the tumour accumulates somatic mutations and evolves to produce multiple genetic subclones (21). The impact of this genetic diversity on tumour immunogenicity is two-fold. An increase in mutation load would result in higher neoantigen expression and increased chances for the neoplasm to be recognized by the immune system. Conversely, the increasing subclonal distribution of neoantigens would direct the immune system to discrete subpopulations of the most immunogenic tumour cells. This in turn would shield less immunogenic subclones from the antitumour attack (45).

In this study, which to our knowledge is the first analysis of neoantigens in MF, we found that the neoantigen load mirrors the mutational load of MF and increases during disease progression. Our experimental approach using microdissected tumour tissue and deep exome sequencing allowed for identification of a markedly higher number of non-synonymous mutations (median 3,217) than previous MF studies (42-102) (27,46,47). The neoantigen load in our MF samples was also much higher than other malignancies known to have a high neoantigen load such as malignant melanoma (median 121) and lung adenocarcinoma (median 335) (48). The differences are not only quantitative, as we were able to detect numerous frameshift mutations (median 2,604) which have hardly been captured in previous studies. Frameshift mutations are an essential source of neoantigens because they often produce highly immunogenic peptides due to global structural aberrations that render the peptide dissimilar from self (49). Thus, MF can be viewed as a neoplasm of high immunogenic potential expressing a significant number (median 328) of high strength neoantigenic peptides.

Analysis of the subclonal heterogeneity of the neoantigens by bioinformatic deconvolution of phylogenetic trees and by multisampling distinct lesions of MF revealed a complex neoantigenic landscape. Our analysis demonstrated that different cutaneous lesions of MF exhibit highly diverse repertoires of non-overlapping neoantigens. The most informative was the analysis of six lesions from a single patient (Figure 6B) which did not share a single antigenic peptide. Similarly, there was overall poor overlap between neoantigens in plaques and tumours from the same patient. Thus, a single patient with MF presenting with numerous skin lesions may be considered as having a collection of multiple, immunologically different neoplasms.

Not only did different lesions vary by their neoantigens but significant neoantigenic heterogeneity was also detected in different lymphoma subclones. Using a bioinformatic approach we were able to show that a large proportion of neoantigens map to the subclones (clades) and that this proportion increased during stage progression. Although it is tempting to speculate that this high proportion of subclonal neoantigens will render advanced stage MF resistant to immunotherapy (22), we have to acknowledge certain limitations of our computational approach. The phylogenetic trees were constructed by statistical modeling of point mutation distributions in the sample and were not verified by single-cell sequencing. Therefore we cannot with certainty equate a branch of the phylogenic tree with a clone of tumour cells.

Although there was a clear increase in the number of neoantigens between early stage plaques and lesions in the late stage disease, it has not escaped our attention that the clinically more advanced lesions of tumours did not have a higher number of antigens (some even had a lower neoantigen load) compared to late stage plaques. This could not have been explained by a lower degree of genetic heterogeneity because the tumours had a highly branched subclonal architecture. We hypothesize that the reduction in neoantigen expression might be a result of immune editing, whereby the cells bearing the most immunogenic neoantigens are negatively selected by the immune system (50).

Previous studies have reported that very few neoantigens are shared across patients in high mutation load malignancies (48) and as already mentioned, our cohort of MF patients did not share any neoantigenic peptides. However, several peptides were commonly found in some patients (Figure 6C) and these could represent potential therapeutic targets. We therefore searched for known homologous immune epitopes of the most frequently observed neoantigens (51). Although none of the homologous sequences were an exact match to our mutant peptides, there were numerous promising partial matches (90% sequence similarity) to immunogenic human sequences and the sequences of human pathogens such as *Mycobacterium tuberculosis* and protozoa (*Leishmania* and *Trypanosoma*) (Table 1). This observation was particularly interesting because neoantigenic peptides homologous to human pathogens are known to be robust activators of the immune response (52)(35). Other notable homologous epitopes included those from proteins implicated in other cancers, such as the ENA family from breast cancer (53), and baculoviral IAP repeat-containing protein 6 from brain cancer (54). Future studies should validate candidate neoantigen expression at the protein level and their ability to elicit T-cell activation.

In conclusion, we have shown a bewildering degree of neoantigen heterogeneity in MF. Among hundreds of detected strong neoantigens there is little overlap between different individuals, between lesions in the same individual and between different subclones within the same lesion. We hypothesize that neoantigen heterogeneity may be an important factor limiting efficacy of immunotherapy in MF, and probably in other highly mutated, genetically heterogeneous cancers. Our results suggest that neoantigenic heterogeneity could serve as a potential biomarker of tumour immunogenicity and response to immunotherapies. Moreover, the observed increase in neoantigen heterogeneity during progression of MF provides rationale for early immunotherapy in this disease.

## Acknowledgments

We extend our gratitude to Dr. Thomas Salopek, Mrs. Rachel Doucet and the nursing staff of Kaye Edmonton Clinic for their assistance with sample collection. We thank Anne-Mette Bjerregaard for her assistance with running MuPeXi. Thanks to Dr. John Elliott, Dr. Andrew Mason and Dr. Gane Wong for their advice on the project. This study was supported by grants from the following sources: Canadian Dermatology Foundation (CDF RES0035718), University Hospital Foundation (University of Alberta), Bispebjerg Hospital (Copenhagen, Denmark), Danish Cancer Society (Kræftens Bekæmpelse R124-A7592 Rp12350) and an unrestricted research grant to R.G. from the Department of Medicine, University of Alberta. A.S. was supported by scholarships from the Canadian Institutes of Health Research (CIHR), Alberta Innovates and the University of Alberta.

## Authors’ contributions

AS contributed to study design, conducted bioinformatics, analyzed the data and wrote the manuscript. DH conducted bioinformatics analysis. AI and SO performed wet lab experiments. PS and GV helped with visual representations of data. RG designed the experiments, supervised data analysis, and edited the manuscript. All authors approved the final version of this manuscript.

## References

1. Korgavkar K, Xiong M, Weinstock M. Changing incidence trends of cutaneous T-cell lymphoma. JAMA Dermatol. 2013;149:1295–9.

2. Iyer A, Hennessey D, O’Keefe S, Patterson J, Wang W, Wong GK-S, et al. Skin colonization by circulating neoplastic clones in cutaneous T-cell lymphoma. Blood. 2019;134:1517–27.

3. Hamrouni A, Fogh H, Zak Z, Ødum N, Gniadecki R. Clonotypic Diversity of the T-cell Receptor Corroborates the Immature Precursor Origin of Cutaneous T-cell Lymphoma. Clin Cancer Res. 2019;25:3104–14.

4. Agar NS, Wedgeworth E, Crichton S, Mitchell TJ, Cox M, Ferreira S, et al. Survival outcomes and prognostic factors in mycosis fungoides/Sézary syndrome: validation of the revised International Society for Cutaneous Lymphomas/European Organisation for Research and Treatment of Cancer staging proposal. J Clin Oncol. 2010;28:4730–9.

5. Trautinger F, Eder J, Assaf C, Bagot M, Cozzio A, Dummer R, et al. European Organisation for Research and Treatment of Cancer consensus recommendations for the treatment of mycosis fungoides/Sézary syndrome - Update 2017. Eur J Cancer. 2017;77:57–74.

6. Sivanand A, Surmanowicz P, Alhusayen R, Hull P, Litvinov IV, Zhou Y, et al. Immunotherapy for Cutaneous T-Cell Lymphoma: Current Landscape and Future Developments. J Cutan Med Surg. 2019;23:537–44.

7. Zackheim HS, Koo J, LeBoit PE, McCalmont TH, Bowman PH, Kashani-Sabet M, et al. Psoriasiform mycosis fungoides with fatal outcome after treatment with cyclosporine. J Am Acad Dermatol. 2002;47:155–7.

8. Sokołowska-Wojdyło M, Barańska-Rybak W, Cegielska A, Trzeciak M, Lugowska-Umer H, Gniadecki R. Atopic dermatitis-like pre-Sézary syndrome: role of immunosuppression. Acta Derm Venereol. 2011;91:574–7.

9. Medrano RFV, Hunger A, Mendonça SA, Barbuto JAM, Strauss BE. Immunomodulatory and antitumor effects of type I interferons and their application in cancer therapy. Oncotarget. 2017;8:71249–84.

10. Chi H, Li C, Zhao FS, Zhang L, Ng TB, Jin G, et al. Anti-tumor Activity of Toll-Like Receptor 7 Agonists. Front Pharmacol. 2017;8.

11. Rook AH, Suchin KR, Kao DM, Yoo EK, Macey WH, DeNardo BJ, et al. Photopheresis: clinical applications and mechanism of action. J Investig Dermatol Symp Proc. 1999;4:85–90.

12. Duarte RF, Schmitz N, Servitje O, Sureda A. Haematopoietic stem cell transplantation for patients with primary cutaneous T-cell lymphoma. Bone Marrow Transplant. 2008;41:597–604.

13. Lesokhin AM, Ansell SM, Armand P, Scott EC, Halwani A, Gutierrez M, et al. Nivolumab in Patients With Relapsed or Refractory Hematologic Malignancy: Preliminary Results of a Phase Ib Study. Journal of Clinical Oncology. 2016;34:2698–704.

14. Khodadoust M, Rook AH, Porcu P, Foss FM, Moskowitz AJ, Shustov AR, et al. Pembrolizumab for Treatment of Relapsed/Refractory Mycosis Fungoides and Sezary Syndrome: Clinical Efficacy in a Citn Multicenter Phase 2 Study. Blood. 2016;128:181–181.

15. Bar-Sela G, Bergman R. Complete regression of mycosis fungoides after ipilimumab therapy for advanced melanoma. JAAD Case Rep. 2015;1:99–100.

16. Sekulic A, Liang WS, Tembe W, Izatt T, Kruglyak S, Kiefer JA, et al. Personalized treatment of Sézary syndrome by targeting a novel CTLA4:CD28 fusion. Mol Genet Genomic Med. 2015;3:130–6.

17. Ansell S, Gutierrez ME, Shipp MA, Gladstone D, Moskowitz A, Borello I, et al. A Phase 1 Study of Nivolumab in Combination with Ipilimumab for Relapsed or Refractory Hematologic Malignancies (CheckMate 039). Blood. 2016;128:183–183.

18. Maier T, Tun-Kyi A, Tassis A, Jungius K-P, Burg G, Dummer R, et al. Vaccination of patients with cutaneous T-cell lymphoma using intranodal injection of autologous tumor-lysate-pulsed dendritic cells. Blood. 2003;102:2338–44.

19. Heinzerling L, Künzi V, Oberholzer PA, Kündig T, Naim H, Dummer R. Oncolytic measles virus in cutaneous T-cell lymphomas mounts antitumor immune responses in vivo and targets interferon-resistant tumor cells. Blood. 2005;106:2287–94.

20. Kim YH, Gratzinger D, Harrison C, Brody JD, Czerwinski DK, Ai WZ, et al. In situ vaccination against mycosis fungoides by intratumoral injection of a TLR9 agonist combined with radiation: a phase 1/2 study. Blood. 2012;119:355–63.

21. Iyer A, Hennessey D, O’Keefe S, Patterson J, Wang W, Wong GK-S, et al. Branched evolution and genomic intratumor heterogeneity in the pathogenesis of cutaneous T-cell lymphoma [Internet]. bioRxiv. 2019 [cited 2019 Oct 19]. page 804351. Available from: https://www.biorxiv.org/content/10.1101/804351v1.abstract

22. McGranahan N, Furness AJS, Rosenthal R, Ramskov S, Lyngaa R, Saini SK, et al. Clonal neoantigens elicit T cell immunoreactivity and sensitivity to immune checkpoint blockade. Science. 2016;351:1463–9.

23. Snyder A, Makarov V, Merghoub T, Yuan J, Zaretsky JM, Desrichard A, et al. Genetic basis for clinical response to CTLA-4 blockade in melanoma. N Engl J Med. 2014;371:2189–99.

24. Rizvi NA, Hellmann MD, Snyder A, Kvistborg P, Makarov V, Havel JJ, et al. Cancer immunology. Mutational landscape determines sensitivity to PD-1 blockade in non-small cell lung cancer. Science. 2015;348:124–8.

25. Wang X, Guo G, Guan H, Yu Y, Lu J, Yu J. Challenges and potential of PD-1/PD-L1 checkpoint blockade immunotherapy for glioblastoma. J Exp Clin Cancer Res. 2019;38:87.

26. Iyer A, Hennessey D, O’Keefe S, Patterson J, Wang W, Salopek T, et al. Clonotypic heterogeneity in cutaneous T-cell lymphoma (mycosis fungoides) revealed by comprehensive whole-exome sequencing. Blood Adv. 2019;3:1175–84.

27. McGirt LY, Jia P, Baerenwald DA, Duszynski RJ, Dahlman KB, Zic JA, et al. Whole-genome sequencing reveals oncogenic mutations in mycosis fungoides. Blood. 2015;126:508–19.

28. Choi J, Goh G, Walradt T, Hong BS, Bunick CG, Chen K, et al. Genomic landscape of cutaneous T cell lymphoma. Nat Genet. 2015;47:1011–9.

29. Van der Auwera GA, Carneiro MO, Hartl C, Poplin R, Del Angel G, Levy-Moonshine A, et al. From FastQ data to high confidence variant calls: the Genome Analysis Toolkit best practices pipeline. Curr Protoc Bioinformatics. 2013;43:11.10.1–11.10.33.

30. Szolek A, Schubert B, Mohr C, Sturm M, Feldhahn M, Kohlbacher O. OptiType: precision HLA typing from next-generation sequencing data. Bioinformatics. 2014;30:3310–6.

31. Bray NL, Pimentel H, Melsted P, Pachter L. Near-optimal probabilistic RNA-seq quantification. Nat Biotechnol. 2016;34:525–7.

32. Bjerregaard A-M, Nielsen M, Hadrup SR, Szallasi Z, Eklund AC. MuPeXI: prediction of neo-epitopes from tumor sequencing data. Cancer Immunol Immunother. 2017;66:1123–30.

33. Jurtz V, Paul S, Andreatta M, Marcatili P, Peters B, Nielsen M. NetMHCpan-4.0: Improved Peptide-MHC Class I Interaction Predictions Integrating Eluted Ligand and Peptide Binding Affinity Data. J Immunol. 2017;199:3360–8.

34. Everaert C, Luypaert M, Maag JLV, Cheng QX, Dinger ME, Hellemans J, et al. Benchmarking of RNA-sequencing analysis workflows using whole-transcriptome RT-qPCR expression data. Sci Rep. 2017;7:1559.

35. Wood MA, Paralkar M, Paralkar MP, Nguyen A, Struck AJ, Ellrott K, et al. Population-level distribution and putative immunogenicity of cancer neoepitopes. BMC Cancer. 2018;18:414.

36. Vita R, Mahajan S, Overton JA, Dhanda SK, Martini S, Cantrell JR, et al. The Immune Epitope Database (IEDB): 2018 update. Nucleic Acids Res. 2019;47:D339–43.

37. UniProt Consortium. UniProt: a worldwide hub of protein knowledge. Nucleic Acids Res. 2019;47:D506–15.

38. Kim S, Scheffler K, Halpern AL, Bekritsky MA, Noh E, Källberg M, et al. Strelka2: fast and accurate calling of germline and somatic variants. Nat Methods. 2018;15:591–4.

39. Ha G, Roth A, Khattra J, Ho J, Yap D, Prentice LM, et al. TITAN: inference of copy number architectures in clonal cell populations from tumor whole-genome sequence data. Genome Res. 2014;24:1881–93.

40. Deshwar AG, Vembu S, Yung CK, Jang GH, Stein L, Morris Q. PhyloWGS: reconstructing subclonal composition and evolution from whole-genome sequencing of tumors. Genome Biol. 2015;16:35.

41. beeswarm: an R package [Internet]. [cited 2019 Dec 13]. Available from: http://www.cbs.dtu.dk/~eklund/beeswarm/

42. Home - GraphPad [Internet]. [cited 2019 Dec 13]. Available from: www.graphpad.com

43. Bardou P, Mariette J, Escudié F, Djemiel C, Klopp C. jvenn: an interactive Venn diagram viewer. BMC Bioinformatics. 2014;15:293.

44. Sterck L. Draw Venn Diagram [Internet]. [cited 2020 Jan 10]. Available from: http://bioinformatics.psb.ugent.be/webtools/Venn/

45. Milo I, Bedora-Faure M, Garcia Z, Thibaut R, Périé L, Shakhar G, et al. The immune system profoundly restricts intratumor genetic heterogeneity. Sci Immunol. 2018;3.

46. da Silva Almeida AC, Abate F, Khiabanian H, Martinez-Escala E, Guitart J, Tensen CP, et al. The mutational landscape of cutaneous T cell lymphoma and Sézary syndrome. Nat Genet. 2015;47:1465–70.

47. Park J, Yang J, Wenzel AT, Ramachandran A, Lee WJ, Daniels JC, et al. Genomic analysis of 220 CTCLs identifies a novel recurrent gain-of-function alteration in RLTPR (p.Q575E). Blood. 2017;130:1430–40.

48. Lee C-H, Yelensky R, Jooss K, Chan TA. Update on Tumor Neoantigens and Their Utility: Why It Is Good to Be Different. Trends Immunol. 2018;39:536–48.

49. Turajlic S, Litchfield K, Xu H, Rosenthal R, McGranahan N, Reading JL, et al. Insertion-and-deletion-derived tumour-specific neoantigens and the immunogenic phenotype: a pan-cancer analysis. Lancet Oncol. 2017;18:1009–21.

50. Yarchoan M, Johnson BA 3rd, Lutz ER, Laheru DA, Jaffee EM. Targeting neoantigens to augment antitumour immunity. Nat Rev Cancer. 2017;17:209–22.

51. Mauriello, Mauriello, Zeuli, Cavalluzzo, Petrizzo, Tornesello, et al. High Somatic Mutation and Neoantigen Burden Do Not Correlate with Decreased Progression-Free Survival in HCC Patients not Undergoing Immunotherapy. Cancers. 2019;11:1824.

52. Rubio-Godoy V, Dutoit V, Zhao Y, Simon R, Guillaume P, Houghten R, et al. Positional scanning-synthetic peptide library-based analysis of self- and pathogen-derived peptide cross-reactivity with tumor-reactive Melan-A-specific CTL. J Immunol. 2002;169:5696–707.

53. Di Modugno F, Bronzi G, Scanlan MJ, Del Bello D, Cascioli S, Venturo I, et al. Human Mena protein, a serex-defined antigen overexpressed in breast cancer eliciting both humoral and CD8+ T-cell immune response. Int J Cancer. 2004;109:909–18.

54. Chen Z, Naito M, Hori S, Mashima T, Yamori T, Tsuruo T. A human IAP-family gene, apollon, expressed in human brain cancer cells. Biochem Biophys Res Commun. 1999;264:847–54.

