## Supplementary File for "The neoantigen landscape of mycosis fungoides"

### Supplementary data

**Table S1:** Characteristics of patients and samples used in the study

| Patient (age, sex) | Sample ID | Lesion type | Diagnosis & stage |
|---|---|---|---|
| MF 4 (69, M) | MF4_1P | Plaque | Mycosis fungoides IIB |
| | MF4_2T | Tumour | |
| | MF4_3P | Plaque | |
| | MF4_4T | Tumour | |
| | MF4_5P | Plaque | |
| | MF4_7T | Tumour | |
| MF9 (42, F) | MF9P | Plaque | Mycosis fungoides IA |
| MF10 (56, M) | MF10P | Plaque | Mycosis fungoides IB |
| MF11 (56, M) | MF11_1P | Plaque | Mycosis fungoides IIB |
| MF15 (65, M) | MF15P | Plaque | Mycosis fungoides IB |
| MF19 (74, M) | MF19_2P | Plaque | |

| | | | |
|---|---|---|---|
| | MF 19_3T | Tumour | Mycosis fungoides IIB |
| MF20 (70, M) | MF20 | Plaque | Mycosis fungoides IB |
| MF27 (71, M) | MF27P | Plaque | Mycosis fungoides IA |
| MF29 (87, F) | MF29_1P | Plaque | Mycosis fungoides IA |
| | MF29_2P | Plaque | |
| MF34 (65, M) | MF34T | Tumour | Mycosis fungoides IIB |
| | MF34_1P | Plaque | |
| | MF34_2T | Tumour | |
| MF36 (64, M) | MF36P | Plaque | Mycosis fungoides IA |
| MF38 (76, M) | MF38_1P | Plaque | Mycosis fungoides IIB |
| | MF38_2T | Tumour | |
| MF40 (59, F) | MF40_1P | Plaque | Mycosis fungoides IIB |
| | MF40_2T | Tumour | |

**Table S2:** Characteristics of CTCL studies used in meta-analysis

| **Study** | **Sample type** | **Sequencing method** | **Sequencing depth (x)** | **Number of samples** |
|---|---|---|---|---|
| Choi et al. | Sézary Syndrome | Whole exome sequencing | Range 142.219-333.623 | 31 |
| McGirt et al. | Mycosis fungoides | Whole genome sequencing | Range 32.04-44.24 | 5 |

**Figure S1: Comparisons of neoantigens from our datasets with those of Choi at al.** (28) **and McGirt et al.** (27) This beeswarm plot shows putative neoantigens prior to filtering. Due to the extensive size of the dataset, a random 1% of all data points were plotted to demonstrate the overall distribution and density of the data. The vertical axis shows mutant peptide binding strength as a percentile rank, with lower values representing increasingly strong binding peptides to HLA types. 0.5% rank (dashed line) represents the commonly used cutoff below which peptides are considered strong enough binders to be neoantigens. The width of each plot represents the quantity of neoantigens at each binding strength. Overall, our dataset had a many fold greater number of putative neoantigens compared to the Choi and McGirt datasets. The slight difference in median binding strength is likely due to the vastly greater size of our dataset (57% rank) compared to the Choi dataset (52% rank) and the McGirt dataset (52% rank). For the McGirt and Choi datasets, there was no RNA data or separation by lesion stages. Consequently, the median filtered neoantigen load was 40-46 per sample (supplementary **Figure S2&S3**)

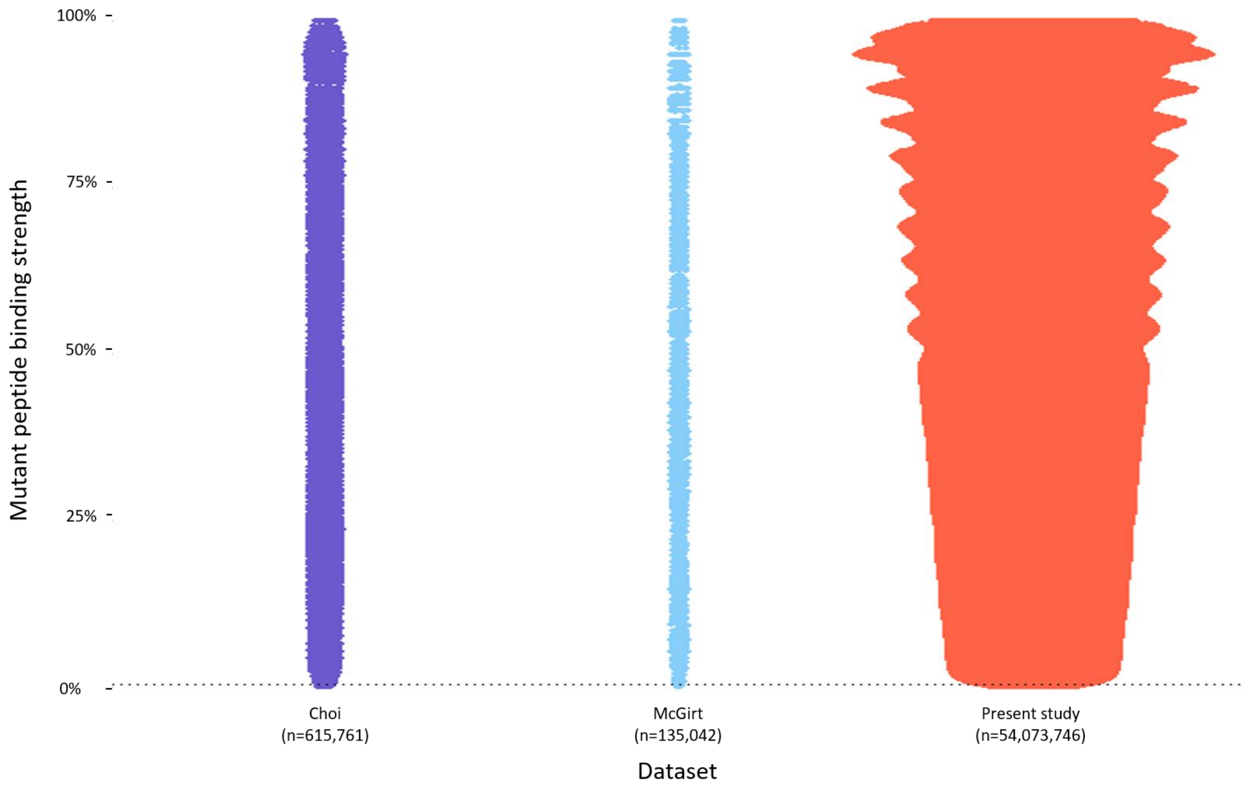

**Figure S2: Characteristics of the dataset from Choi et al.** (28)**. A:Tumour mutation burden.** Samples are arranged in descending order of TMB. Missense mutations comprise 96% of the non-synonymous mutations. **B: Filtered neoantigen load.** All filters were applied with the exception of the RNA filter as expression data was not available. The median number of filtered neoantigens per sample was 40.

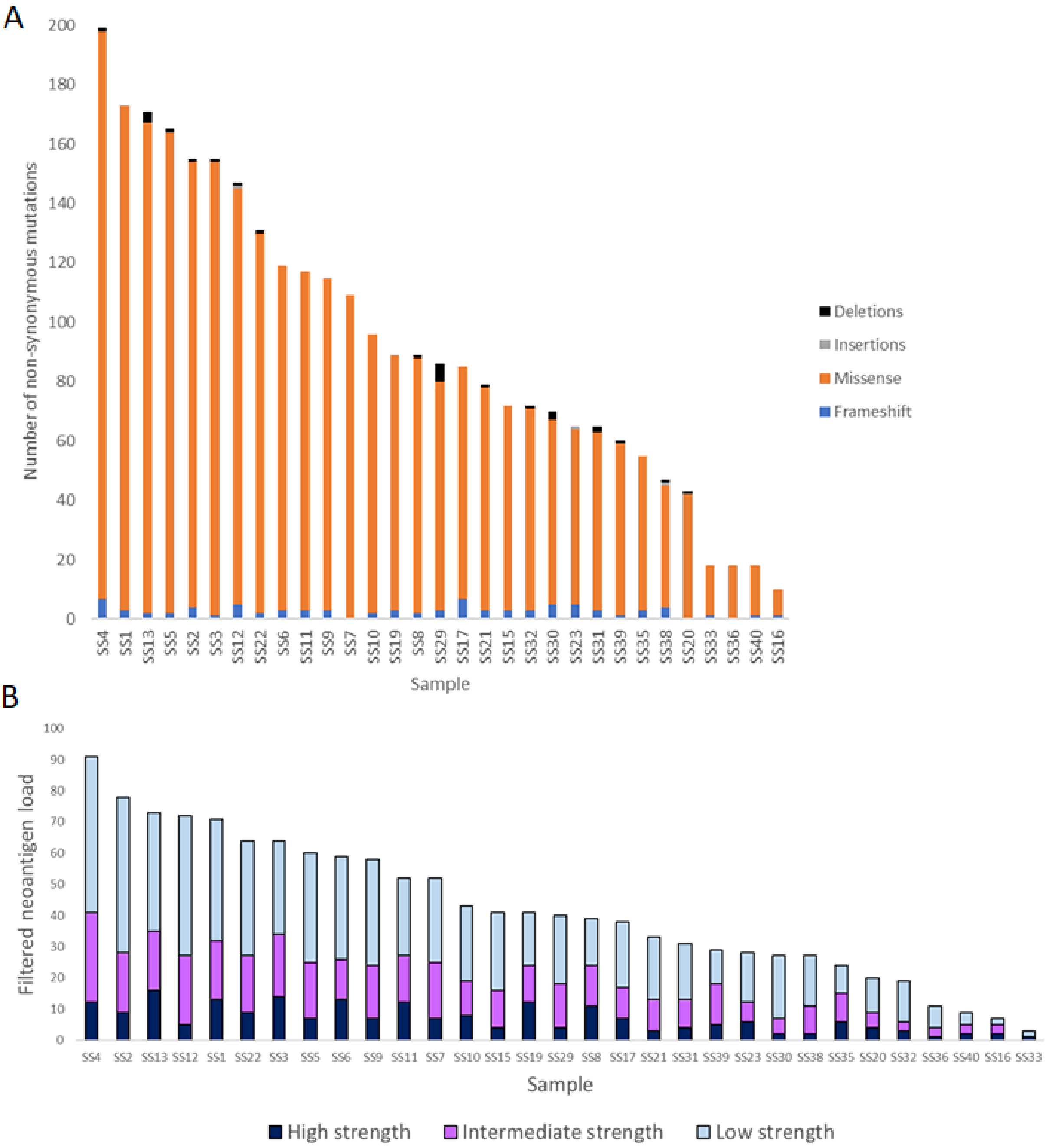

**Figure S3: Characteristics of the dataset from McGirt et al.** (27)**. A:Tumour mutation burden.** Samples are arranged in descending order of TMB. Missense mutations comprise 98% of the non-synonymous mutations. **B: Filtered neoantigen load.** All filters were applied with the exception of the RNA filter as expression data was not available. The median number of filtered neoantigens per sample was 46.

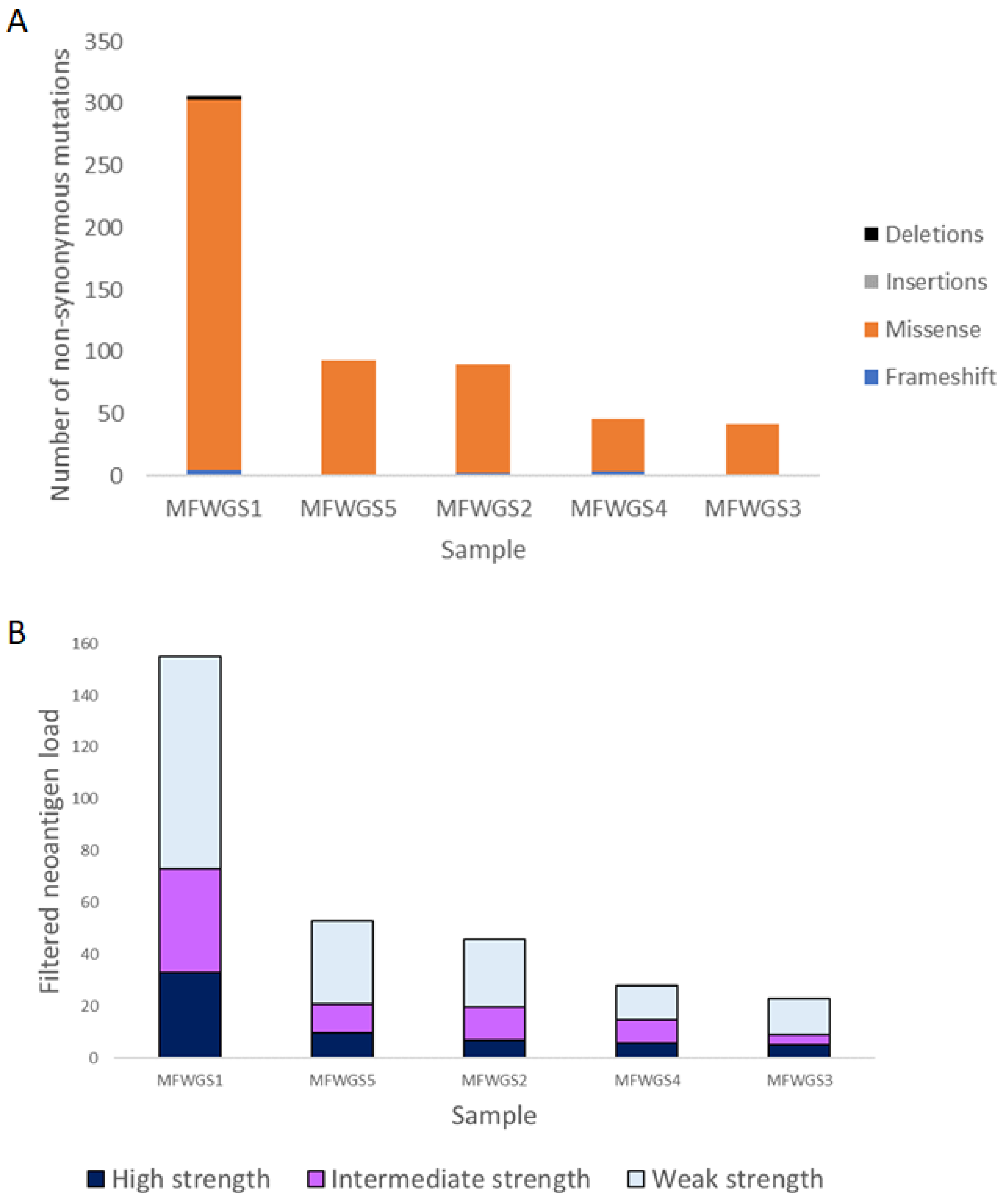
